# Tiberius: End-to-End Deep Learning with an HMM for Gene Prediction

**DOI:** 10.1101/2024.07.21.604459

**Authors:** Lars Gabriel, Felix Becker, Katharina J. Hoff, Mario Stanke

## Abstract

**Motivation:** For more than 25 years, learning-based eukaryotic gene predictors were driven by hidden Markov models (HMMs), which were directly inputted a DNA sequence. Recently, Holst et al. demonstrated with their program Helixer that the accuracy of *ab initio* eukaryotic gene prediction can be improved by combining deep learning layers with a separate HMM postprocessor.

**Results:** We present Tiberius, a novel deep learning-based *ab initio* gene predictor that end-to-end integrates convolutional and long short-term memory layers with a differentiable HMM layer. Tiberius uses a custom gene prediction loss and was trained for prediction in mammalian genomes and evaluated on human and two other genomes. It significantly outperforms existing *ab initio* methods, achieving F1-scores of 62% at gene level for the human genome, compared to 21% for the next best *ab initio* method. In *de novo* mode, Tiberius predicts the exon-intron structure of two out of three human genes without error. Remarkably, even Tiberius’s *ab initio* accuracy matches that of BRAKER3, which uses RNA-seq data and a protein database. Tiberius’s highly parallelized model is the fastest state-of-the-art gene prediction method, processing the human genome in under 2 hours.

**Availability and Implementation:** https://github.com/Gaius-Augustus/Tiberius

**Contact:** {}

## Introduction

Gene prediction is the task of finding genes and their exon-intron structure in a genome and is a fundamental step in the analysis of a newly sequenced genome. Its output includes the coordinates of exons, the set of protein sequences encoded by the genome and is the basis for most downstream analysis tasks. In eukaryotes, the gene prediction task has not yet been solved with an accuracy that is satisfactory for most tasks. In particular, comparative studies that correlate differences in observed phenotypes with differences in genes require high genome annotation accuracy [25]. Plans to sequence the genomes of all known eukaryotic species suggest a demand of more than a million genomes that require annotation in the next decade [20].

Most currently deployed pipelines for gene prediction use evidence from previously identified proteins and their accuracy also benefits from integrating RNA-seq data [10, 7, 14]. Furthermore, RNA-seq is widely recognized to be required for highly accurate predictions [18]. However, taking mammals as an example, currently there are 341 mammalian species for which a genome assembly is deposited at the National Center for Biotechnology Information (NCBI) but no RNA-seq available (48% of the total). If a similar genome annotation quality could be reached without RNA-seq, a substantial amount of time and other resources could be saved.

Many existing genome annotation pipelines, such as BRAKER [5, 10] or NCBI’s Eukaryotic Genome Annotation Pipeline [29] already have machine learning or statistical components, such as Markov chains or splice site pattern recognizers that exploit patterns in the genome itself whose parameters need to be trained for a given clade. If *only* evidence from the target genome is used, this is referred to as *ab initio* gene prediction. If also evolutionary evidence from multiple (aligned) genomes is used, this is referred to as *de novo* gene prediction. Formally, the *ab initio* gene prediction task is to give each position in the input genome sequence *S* ∈ {A, C, G, T}^*T*^ a biological label of some set of labels *Q* that includes exons, introns and intergenic region, i.e. to produce an output *Y* ∈ *Q*^*T*^. This formulation is a *simplification* because it does not account for alternative splicing and overlapping and nested genes. Nevertheless, this simplified task is difficult and at the core of genome annotation.

A hidden Markov model (HMM) is a generative probabilistic model *P* (*X, Y*) of an observed sequence *X* = *f* (*S*), where *f* can be any transformation of the genomic input to the HMM input, and a “hidden” sequence *Y*. Traditional methods *directly* use the DNA sequence as HMM input (i.e. *X* = *S*). Such HMMs with direct DNA input and slight generalizations thereof have been used for gene prediction for more than 25 years [12, 27]. Arguably the majority of genomes is currently annotated using annotation pipelines that employ HMMs such as the NCBI pipeline which uses the “GenScan”-like HMM Gnomon [29] and BRAKER, which uses AUGUSTUS [27] and GeneMark [21].

Recently, gradient-based training of deep learning models has been adapted for gene prediction. Stiehler *et al*. [28] introduced a deep neural network that uses a combination of convolutional neural network (CNN) and long short-term memory (LSTM) layers. Their program Helixer is a sequence-to-sequence model *f* that outputs a matrix *X* ∈ [0, 1]^*T ×*|*Q*|^ such that *X*[*i, q*] is the estimated probability that the *i*-th genome position has biological label *q*. However, Helixer did not yet predict gene structures. To provide gene predictions, Holst *et al*. added a HMM-based ‘post-processor’ that uses the Viterbi algorithm with input *X* = *f* (*S*) to infer a most likely gene structure 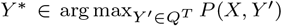 This version of Helixer that is post-processed by an HMM is here referred to as just ‘Helixer’. It was hitherto arguably the overall most accurate *ab initio* gene predictor, although Holst *et al*. did not evaluate Helixer with regard to the usual measures of gene prediction accuracy, such as recall and precision for the prediction of exons and genes. With it, Holst et al. have shown that deep learning has the potential to push the limits of the ‘shallow’ learning approaches of classical HMMs such as AUGUSTUS. However, in the Helixer model, *f* is not adapted to the HMM that generates gene structures. Instead, the model *f* is trained in an intermediate sequence classification task without inductive bias of the true biological limitations of a gene structure. All biological constraints imposed on a full gene structure have to be learned and stored in the parameters of *f*.

The accurate prediction of a gene structure is highly dependent on the correct identification of state transitions, such as the borders of exons. Holst *et al*. have shown that a general-purpose deep learning sequence-to-sequence model can achieve high base-level precision, but struggles to precisely locate exon boundaries, which could lead to low exon- and gene-level accuracy. Marin *et al*. compared DNA language models with classical HMM-based AUGUSTUS for human gene finding. the best performing DNA language model, Nucleotide Transformer [8], did “not approach the highly specialized AUGUSTUS”. Marin *et al*. concluded that “more specialized downstream models are still needed to accurately predict gene structure”.

Recently, Becker and Stanke developed a *HMM layer* [3] in the context of multiple sequence alignments with profileHMMs. This HMM layer is a special case of a recurrent neural network that can be used in both supervised and unsupervised settings jointly with other layers. We utilize the HMM layer in a model inspired by Helixer, incorporating several enhancements. Instead of relying on a post-processor, we integrate the HMM during *both* training and inference. Therefore, we leverage end-to-end training with the full inductive bias of a gene structure (consistent reading frame, start- and stop codons and splice site patterns). In training mode, the output of our full model with HMM are the posterior state probabilities Ŷ[*i, q*] = *P* (*Y*_*i*_ = *q*|*X*). We train it by minimizing a misclassification loss *L*(Ŷ, *Y*_*true*_). In contrast, Helixer minimizes a loss function *L*(*X, Y*_*true*_) that only depends on *X* and thus is independent of the HMM.

Our new gene finder Tiberius outperforms Helixer on *ab initio* mammalian genome annotation (gene-level F1-score 55% versus 19%) runs significantly faster on GPU than Helixer as well as established CPU-based tools and can be highly parallelized. We also compared Tiberius with BRAKER3, as a representative of current state-of-the-art pipelines that integrate RNA-seq and protein sequence data. Despite Tiberius not using extrinsic evidence, it performed slightly better in our test setting that disregarded alternative splicing.

## Methods

### Dataset

A dataset comprising assemblies of 37 mammalian species (see Table 1) was used for training and evaluation of Tiberius. Genomes were soft-masked using RepeatModeler2 [9], RepeatMasker (http://www.repeatmasker.org) and Tandem Repeats Finder [4]. RefSeq reference annotations [29] were retrieved from NCBI. For details, we refer to the Supplementary Methods. This dataset includes model species such as human and mouse, covering a broad phylogenetic range of mammalian species (see Supplementary Figure1). The gene prediction challenge here arises from a relatively low gene density, large numbers of exons per gene, GC-content heterogeneity, a large genome size, and many long introns. For example, in human, about half of the genes have at least one intron of size 10Kb or larger. Dataset properties are given in Supplementary Tables S1 and S2. To filter out annotations that suffer from apparent false negatives, we included only species in the dataset with a BUSCO [22] completeness greater than 90%.

**Table 1.** Average runtime (h:m) of the five gene prediction tools used in the benchmarking experiments.

| A100 GPU |  | 48 threads CPU |  |  |  |
| --- | --- | --- | --- | --- | --- |
| Tiberius | Helixer | AUGUSTUS | BRAKER3 | GALBA | BRAKER2 |
| **1:39** | 8:54 | 2:25 | 48:53 | 35:12 | 15:36 |

As test species, we chose the mammal with the arguably best annotated genome, human (*Homo sapiens*), as well as two others for diversity: cow (*Bos taurus*) and beluga whale (*Delphinapterus leucas*). The training and validation genomes were controlled for phylogenetic proximity to the test species (Supplementary Figure 1). We did not include any species from the taxonomic groups of *Hominidae, Ruminantia*, and *Cetacea* in the test or validation set. The rationale is that the performance on the test species shall be an estimate of the performance of Tiberius on other mammalian genomes. The minimal evolutionary distance from a training species to human is 43 MYA and to cow and beluga whale 64 MYA.

For the validation of the Tiberius model training, *Panthera pardus* and *Rattus norvegicus* were randomly selected from the training species set. Their genomic data were subsequently excluded from the training process and only used for hyperparameter selection.

For each gene within the reference annotations, the transcript with the longest coding sequence was chosen to generate unambiguous training labels. This is because the Tiberius model’s output is designed to assign a single label per base position, excluding the possibility of representing alternative splicing isoforms.

### Tiberius’ Architecture

The architecture of the Tiberius model integrates CNNs, LSTMs and a differentiable and parallelized HMM layer (Figure 1). Although the pre-HMM architecture shares similarities with the Helixer model introduced by Holst *et al*., the architecture of Tiberius distinguishes itself through several unique features: an HMM layer, repeat information as an additional input, training loss, residual connection, number of output class labels and a larger parameter count.

**Fig. 1:**
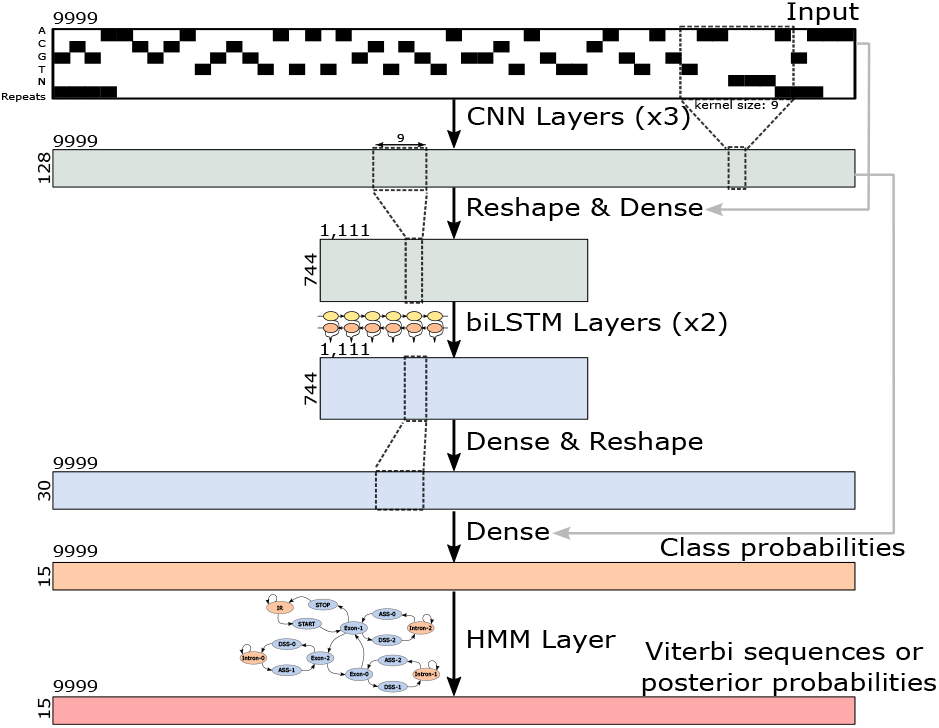
Illustration of the CNN-LSTM architecture of the Tiberius model for gene structure classification at each base position. The HMM layer computes posterior probabilities or complete gene structures (Viterbi sequences). The model has approximately eight million trainable parameters and it was trained with sequences of length *T* = 9999 and a length of *T* = 500,004 was used for inference.

The input to the model is a sequence representing the one-hot encoded nucleotide sequence over the alphabet {A, C, G, T, N} stacked with a track of masked repeat regions. The model outputs, during training, probabilities for 15 gene structure classes we define below and, during inference, label sequences.

The model is capable of processing sequences of variable lengths. This allows it to be trained on shorter sequences to reduce training time and subsequently to be evaluated on longer sequences in order to get a longer context length. The architecture is designed to predict one strand at a time. To make a prediction on the reverse strand, Tiberius inputs the reverse complement of the input sequence into the model. This allows it in principle to predict overlapping or nested genes on opposite strands (see Supplementary Fig. 5 for an example).

### Tiberius Training

The Tiberius model was trained using batches of seamless tiles of length 9999. The training process took 15 days on a machine equipped with four A100 GPUs (80GB memory each). During a first phase, the pre-HMM training, we trained the model without the HMM layer for six days. Afterwards, in phase two, we fine-tuned the model end-to-end for nine days, including the HMM layer. The software documentation provides instructions that can be adapted for a different training set. We used the Adam optimizer [15] with a learning rate of 10^−4^ throughout training. The batch size per GPU was 250 during the pre-HMM training and 128 during the fine-tuning.

For validation, we evaluated the model’s performance after every 1000 training steps using selected genome segments from the validation species,leopard and rat. The evaluation consisted of making full gene structure predictions for these validation data. Training was terminated when no improvement in gene and exon level accuracies was observed on the validation set. We selected the final model based on the highest combined F1-scores for validation exon and gene level accuracies.

### F1-Loss

A notable challenge for training a gene prediction model with genomic data is the underrepresentation of exon classes in genomic data.

For instance, on the forward strand of the human genome, coding exon regions constitute only ∼1% of the genomic sequence, while ∼16% are intron regions and ∼83% intergenic regions. The standard cross-entropy loss alone is not a suitable representation of how well the model predicts the minority class (exons). The precision and recall on gene and exon level are highly sensitive to changes in the prediction of individual labels and require precise identification of exon boundaries.

Loss functions with class weights typically fail to optimize both precision and recall simultaneously, particularly for underrepresented classes such as exons, as they are designed to minimize overall error rates but do not effectively manage the trade-offs between false positives and false negatives (here: exons) [30]. To address this imbalance and improve model performance on gene prediction metrics, we introduce the F1-loss function. This function creates a loss function based on the estimated F1-score for the exon class labels, which we use in combination with the categorical cross-entropy (CCE) loss. The F1-loss aims to improve the model’s accuracy in predicting exon labels, and it aligns the training objectives better with the evaluation metrics (at least on base level). The F1-loss is defined as

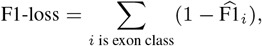

where 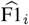 is the estimated F1-score of class *i*. We describe the computation of 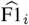 in the Supplementary Methods.

For training the Tiberius model, we use the F1-loss in combination with the CCE-loss function. The latter takes all class labels into account. In instances where a sequence has no exon labels, the estimated false positive rate 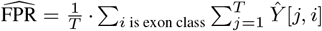 is calculated instead of the F1-loss. The CCE-F1-loss is then computed as:

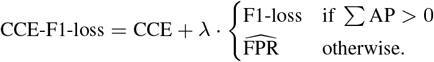

In our experiments we chose *λ* = 2. The CCE-F1-loss is computed for each sequence in a batch independently and the per-sequence losses are averaged.

#### Ablations

To assess the impact of key features in our model, we conducted ablation studies on Tiberius. Each ablation modified a specific feature of the original model and the ablated model was trained in the same manner as the pre-HMM training of Tiberius (see Tiberius Training):

- Tiberius_preHMM: The model without the HMM layer.
- Tiberius_no_sm: Removed the repeat masking track from input.
- Tiberius_CCE: Used CCE-loss, instead of CCE-F1-loss.
- Tiberius_small: Reduced the number of parameters (to approximately two million) by halving the filter and unit sizes of the CNN-/biLSTM layers.
- Tiberius_5class: Removed splice site classes, by mapping the 15 output classes to a set of five labels (IR, Intron, Exon-0, Exon-1, Exon-2) more similar to the one used by Helixer before loss calculation.
- Tiberius_5class_CCE: Used above five labels and the CCE loss.

### HMM Layer

HMMs, traditionally used in gene finding, can be designed to enforce biological constraints, e.g. reading-frame consistency. A most probable sequence of states for an input sequence can be computed with the Viterbi algorithm. Following Holst *et al*., we decode complete gene structures from an input matrix *X* of position-wise class probabilities.

We developed a novel HMM layer [3] using TensorFlow, which is an integral component of the model during fine-tuning and inference. It supports all functionality of a traditional deep learning layer: it is vectorized (i.e. it runs on a batch of independent sequences in parallel), it runs on GPU and it is differentiable (i.e. it can be used for end-to-end fine-tuning). It also supports a parallel mode that is able to parallelize common HMM algorithms over segments of the input sequences, as described later, in order to maximize utility of the available GPUs.

#### HMM architecture

We use an HMM with 15 hidden states of which 11 states are for exon positions, three for intron positions and one for intergenic positions (Figure 2). The exon states are further divided into three states corresponding to the different reading frames and nine states that are specifical for exon border positions. The border states are two states for acceptor and donor splice sites for transitions to and from the intron states and two states for start and stop codons of a gene.

**Fig. 2:**
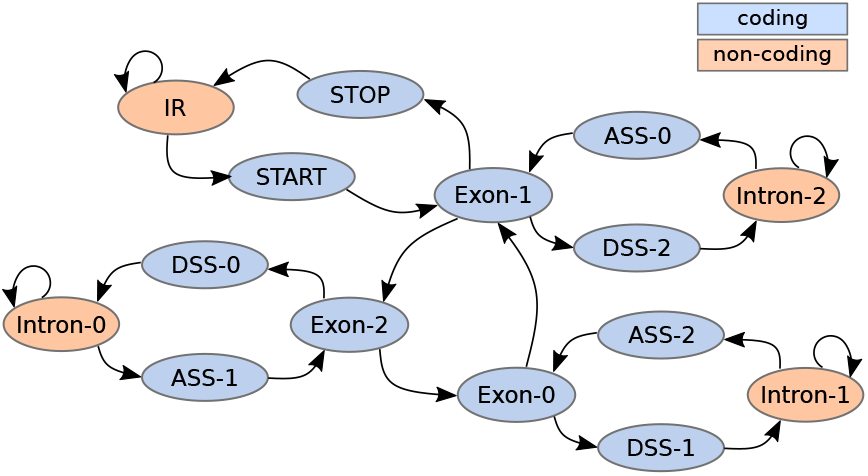
The states of the HMM used for inference with Tiberius and the transititons between them. The 11 blue states correspond to coding-exon positions, each subdivided by reading frame i: Exon-i represents non-border positions within an exon, while ASS-i (acceptor splice site) and DSS-i (donor splice site) states are the first and last position of an exon that starts and ends with reading frame *i*, respectively. The four red states represent a non-coding positions: intergenic region (IR) or intron.

The inputs to our HMM are position-wise class probabilities and sequences of right- and left-adjusted, overlapping triplets. The 15 input classes correspond one-to-one to the states of the HMM. The HMM is designed to enforce canonical splice site patterns, start and stop codons, while also preventing in-frame stop codons within exons. For details, we refer to the Supplementary Methods. Note that the design does not prevent in-frame stop codons that are spliced across two exons, and these may appear in the predicted gene structures; such genes are currently filtered out during post-processing. Allowed splice site patterns and their relative weighting can in principle be specified by the user or even be learned. We benefit from the small number of states of Tiberius’ HMM in the parallel implementation of the Viterbi algorithm.

#### Fine-tuning with HMM

The HMM has only 24 parameters that were not trained. The 23 transition parameters (one for each edge in Figure 2) are determined using expected lengths for intergenic, intron, and exon regions. The emission distribution maps input classes directly to corresponding states and has only a single smoothing parameter to account for incorrect inputs. See Supplementary Methods Section 3 for more details. Although the HMM itself is not trained, the gradient of the loss computed from the HMM’s output with respect to the HMM’s input was computed, and the HMM serves during training as an inductive bias for biological constraints of the gene structures.

#### Parallel Viterbi

We found that for long inputs the HMM layer is the computational bottleneck. The Viterbi algorithm requires to compute dynamic programming variables sequentially for each time step that depend on the variables of the previous step. This is problematic for long sequences. Note that the LSTM is also sequential but receives, like in the Helixer model, an input that is shortened by a reshaping operation. To solve this issue, we employ a parallel variant of Viterbi, which can run in parallel on segments of a sequence. This can be done by conditioning on the states at the segment borders and computing *local* Viterbi variables under these conditions. An equivalent result to the sequential Viterbi can be computed from these local variables in another *global* pass, this time requiring only one step *per segment*. More details are given in the Supplementary Methods. This procedure trades off an increase in the total number of necessary computations proportional to the number of HMM states with the possibility of high parallelization. The parallel Viterbi algorithm greatly improves the speed of Tiberius and increases its GPU utilization while being functionally equivalent. With parallelization it runs more than 17 times faster on GPU (Supplementary Table 5).

### Inference of Gene Structures

The Tiberius model employs the HMM layer and the parallel Viterbi algorithm to infer complete gene structures. Let *S* denote the input genome stacked with the corresponding soft masking track. This sequence is partitioned into seamless tiles {*S*_1_, *S*_2_, …, *S*_*n*_}, where each *S*_*i*_ has length *T* (divisible by 9 for implementation reasons, see above). For our experiments, we compared different tile lengths for inference on the validation species (Supplementary Table 8) and selected a tile length of *T* = 500,004, approximately 50 times the length used during training In order to perform the inference, the Tiberius model is divided into two main components, which are functions that are composed as 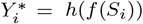. Here, *h* executes the Viterbi algorithm of the HMM layer and *f* is the function computed by the preceding CNN-LSTM layers. Initially, the CNN-LSTM layers process all tiles to generate preliminary predictions *X*_*i*_ = *f* (*S*_*i*_) ∈ [0, 1]^*T ×*|*Q*|^. To save computational time, a 200-base-long sliding window average is computed along each sequence of predictions for the sum of the exon classes. If the maximum of this average is below 0.8 for a tile *X*_*i*_, the HMM layer is not applied on this tile and no gene is predicted within this tile. For the remaining tiles, *h* is applied to infer the most likely sequence of hidden states 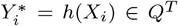 from the HMM, given the predictions *X*_*i*_ by the CNN-LSTM model.

We want to predict gene structures that are fully contained (from start to stop codon state) in *a single* Viterbi parse 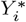. If the hidden state of a Viterbi parse 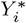 at a tile border is not the intergenic region, the two adjacent tiles are concatenated for a second round of predictions, e.g. computed as 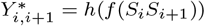. Subsequently, gene structures inferred from 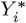 and 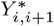 are merged (see Supplementary Methods Section 3). At the end, any gene structures with a coding length smaller than 201 and with in-frame stop codons are removed. The remaining structures are output to a GTF formatted file.

### Tiberius *de novo*

Tiberius has a *de novo* mode that uses evolutionary evidence from multiple unannotated genomes. This evidence, generated by ClaMSA [24] using 64 species in the Zoonomia project alignment [1], provides site-specific coding probabilities. These are summarized into four values per position, representing the logit values for codon start sites. These values are integrated into Tiberius by stacking them to each position’s input data. For training and benchmarking, ClaMSA data were generated using *Mus musculus* and *Homo sapiens* as reference species, respectively. We only include *de novo* results for human as our multiple genome alignment for inference was human-referenced. See Supplementary Methods for more details.

### Accuracy Metrics

To assess the accuracy of predicted gene structures, we adopted metrics commonly used in evaluating gene prediction tools. We compared the predicted gene structures to a reference annotation at two different levels: exon and gene. For both levels, we quantified the number of true positives (TP), false positives (FP), and false negatives (FN). We used the following metrics: *Recall* = TP/(TP+FN) - the proportion of instances correctly identified from the reference, *Precision* = TP/(TP+FP) - the proportion of correct predictions, and the *F1-score* - their harmonic mean. Note that in the field of gene finding methods and their evaluation, the terms ‘specificity’ and ‘sensitivity’ are sometimes used interchangeably with our definitions of ‘precision’ and ‘recall’, respectively. Additionally, we computed the BUSCO completeness, as this is a commonly used metric for evaluating novel genome annotations.

### Benchmarking

Whole-genome predictions were generated for the test species — *Homo sapiens, Bos taurus*, and *Delphinapterus leucas* — using Tiberius and its ablated variants. We compared Tiberius with the other *ab initio* gene finders Helixer and AUGUSTUS, which can use the same input data. Helixer is a very closely related method, as it has a similar architecture and is the only other gene prediction tool based on deep learning. In contrast, AUGUSTUS is a well-established gene finding model which employs a HMM and does not utilize deep learning. For AUGUSTUS, the standard human parameters were used, that were trained in 2010 on 1784 human genes. Helixer was executed with publicly available weights for vertebrate genomes using recommended settings. The three species we used for testing were all included in the list of 315 vertebrate genomes on which Helixer was trained.

In a second set of experiments, we compared Tiberius with the genome annotation pipelines BRAKER2 [5], GALBA [6], and BRAKER3 [10], which utilize extrinsic evidence. In particular, BRAKER2 and GALBA incorporate protein sequences. BRAKER2 uses a large protein database that may include distantly related species. GALBA relies on sequences from a few closely related species. BRAKER3, arguably the most accurate automated genome annotation pipeline currently available [10], incorporates both RNA-seq data and a large protein database as additional inputs. In our experiments, we used the vertebrate partition of OrthoDB v.11 as protein database for BRAKER2 and BRAKER3, excluding – just as for training Tiberius – sequences from either *Hominidae, Ruminantia*, or *Cetacea*, depending on the target species. Additionally, BRAKER3 used ten paired-end short-read RNA-seq libraries for each test species (Table 3). For the GALBA experiments, we selected protein sequences from three closely related species from the Tiberius training set for each test (see Supplementary Methods 3). Unlike Tiberius, these pipelines can report alternative splice forms for each gene. To ensure a fair comparison at the gene level, where additional isoforms can only increase accuracy, we used only the isoform with the longest coding sequence for each gene.

## Results

### Benchmarking Results

The benchmarking experiments assessed Tiberius through comparisons with two distinct groups of gene prediction methods. The first comparison group consisted of other *ab initio* methods - AUGUSTUS, and Helixer - which rely solely on genomic data without extrinsic evidence. The second group, consisting of BRAKER2, BRAKER3, and GALBA, utilized extrinsic evidence for their gene predictions.

#### Comparison of *Ab Initio* Methods

Tiberius consistently outperformed AUGUSTUS and Helixer across all species and metrics (Figure 3). On average, Tiberius achieved an F1-score of 89.7% at exon level and 55.1% at gene level, followed by Helixer (72.9% and 19.3%, respectively) and then AUGUSTUS (67.3% and 12.4%, respectively) (Supplementary Table S6). This trend was evident across all tested species, the highest leading margin for Tiberius was observed in *Homo sapiens*, where Tiberius has 41.8 percent points higher precision and 40.5 percent points higher recall than Helixer on gene level. As an additional performance metric for genome annotation, protein-level BUSCO completeness was calculated for all gene predictions. Tiberius led this metric as well, achieving an average of 96.0% BUSCO completeness for the test species, followed by Helixer at 92.1% and AUGUSTUS at 74.2% (Supplementary Fig. 2).

**Fig. 3:**
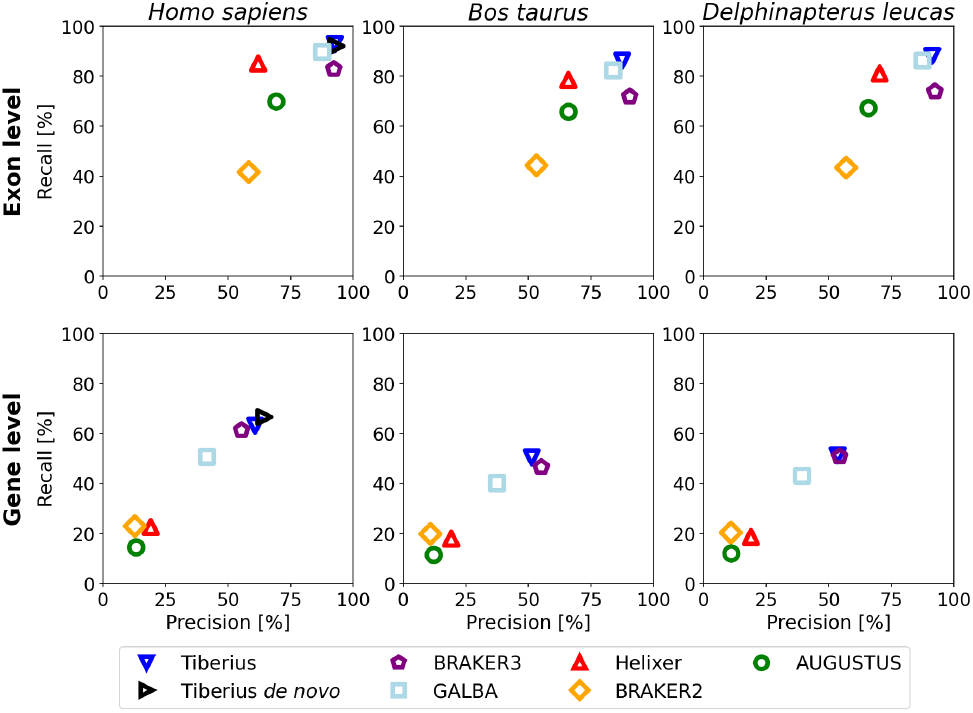
Gene and exon level precision and recall for Tiberius, BRAKER3, GALBA, Helixer, BRAKER2, and AUGUSTUS. Tiberius, Helixer, and AUGUSTUS performed *ab initio* predictions while the other methods additionally incorporated extrinsic evidence: GALBA proteins from related species, BRAKER2 a large protein database, and BRAKER3 a large protein database and RNA-seq. For the human genome, Tiberius was also run *de novo*.

The models of Helixer and Tiberius share many similarities, but their large test accuracy differences may be attributed to several design or implementation choices; these include model size, training loss, output class labels and end-to-end learning with an HMM layer. The most significant impact on training accuracy came from introducing the CCE-F1-loss combined with output labels that include different classes based on reading frame and exon borders (see Ablation Studies and Generalizability). In contrast, Helixer used a weighted CCE-loss without separate exon border classes and uses higher weights for the boundaries of reference exons only [13]. Integrating an HMM layer for end-to-end training further improved test accuracy, increasing F1-scores by 2.6 and 1.1 percentage points at gene and exon levels, respectively. Increasing parameters from 2 million (comparable to Helixer’s 3 million) to 8 million yielded additional improvements of 4.9 and 1.2 percentage points in gene and exon level F1-scores, respectively. Tiberius demonstrates the ability to accurately predict even genes that span large genomic regions. The longest human gene correctly predicted by Tiberius is 328,931 bases long.

#### Comparison with Methods Using Extrinsic Evidence

The integration of protein and RNA-seq data into genome annotation methods has so far been regarded as a significant advantage over *ab initio* predictions. The very recently published BRAKER3, which uses both RNA-seq and protein evidence, represents the current state of the art for genome annotation. For our accuracy comparisons, we eliminated the influence of the prediction of alternative splice forms by using for each gene set, predicted or reference, only the transcript variant with the longest coding sequence.

Compared to state-of-the-art methods that incorporate extrinsic evidence, the *ab initio* predictions by Tiberius match or surpass the performances of the other tools, despite its disadvantage (Figure 3). At gene and exon level, Tiberius achieves an F1-score of 55.1% and 89.7%, which is 1.4% and 6.5% points higher than that of BRAKER3 (53.7% and 83.2%), surpassing also the F1-scores of both GALBA (41.8% and 86.2%) and BRAKER2 (14.9% and 48.7%) (Supplementary Table 6). The increase in accuracy is particularly notable in the case of *Homo sapiens*, where Tiberius surpasses the other methods in all metrics. Tiberius executed in *de novo* mode on human has even higher accuracy on gene level, reaching an F1-score of 65.5% and 92.6% on gene and exon level. In terms of BUSCO completeness, Tiberius achieves the highest completeness for each species, most complete in human with 98.9%.

As illustrated in Supplementary Fig. 6, the integration of extrinsic evidence has the potential to reduce Tiberius’ errors and to further improve gene prediction accuracy. The evolution of other *ab initio* methods, that have incorporated extrinsic evidence, suggests that such integration could ultimately lead to substantial accuracy improvements [33].

### Ablation Studies and Generalizability

We evaluated the impact of key features of Tiberius by retraining ablation models and comparing them to the original model. To reduce computational time, all models in this evaluation, including the default model (Tiberius_preHMM), were trained without the HMM layer. The HMM layer was used only during inference.

#### Softmasking

Retraining Tiberius without the softmasking input track (Tiberius_no_sm) shows a slight decline in prediction accuracy (Supplementary Table S7). The average F1-score was decreased by just 0.2% points on exon level and 0.9% points on gene level. For more distant species, the differences are larger, and Tiberius’s accuracy benefits substantially from the softmasking input (Supplementary Fig. 3). This effect is presumably due to these species having different repeat families that Tiberius_no_sm has not learned.

The minor gains in mammalian species must be weighed against the additional preprocessing step required to mask the repeats before running Tiberius. The Tiberius software provides both options, allowing users to input genomic sequences with or without softmasking.

#### Class Labels and Loss Function

The F1 part of the CCE-F1-loss weights all exon classes equally, regardless of their frequency of occurrence, which is particularly relevant for the positions of the splice sites. Using separate classes for exon borders, as implemented in the 15 classes of the default model, aligns the loss function with the target metrics of exon and gene level accuracies. The loss function and class labels are beneficial because small differences in exon border positions can have a significant impact, even if the majority of an exon segment is predicted correctly.

Our ablation studies demonstrate that both the CCE-F1-loss and separate exon border classes lead to substantial improvements, particularly on gene level accuracy. Compared to Tiberius_preHMM, Tiberius_CCE decreases gene and exon level F1-scores by 8.8 and 2.1 percentage points, while Tiberius_5class decreases them by 11.8 and 5.4. Removing both features causes even a decrease by 21.1 and 10.3 percentage points in gene and exon level F1-scores.

#### Generalizability

To assess Tiberius’s ability to generalize beyond the mammalian training set, we evaluated its performance on species across varying evolutionary distances (Figure 4). The results show robust performance on mammalian test species and a steady decline in accuracy as phylogenetic distance increases. Notably, Tiberius maintains strong performance on more distant vertebrates such as *Gallus gallus* and, at least on exon level, *Danio rerio*. The most divergent species tested, *Populus trichocarpa* and *Solanum lycopersicum*, show the lowest F1-scores, as expected given their large evolutionary distance from the training data. This demonstrates Tiberius’s potential for applicability across domains that are adjacent to mammals, while showing the need for re-training for other clades.

**Fig. 4:**
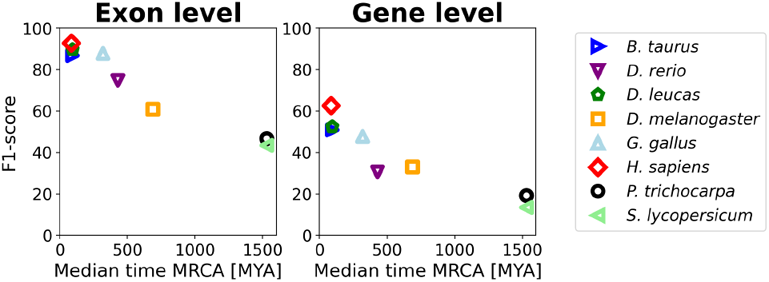
Tiberius accuracy for test species, including non-mammalian species, plotted against the median time from the most recent common ancestor (MRCA) with *Mus musculus*, generated with TimeTree [16].

### Runtime

We ran our experiments for Helixer and Tiberius on a high-performance computing cluster using an NVIDIA A100 GPU with 80Gb memory and 48 CPU threads, but Tiberius makes little use of the CPU cores. BRAKER3, GALBA, BRAKER2, and AUGUSTUS do not support GPU execution and were executed on a node using 48 CPU threads. Among the models tested, the *ab initio* methods had significantly lower runtimes, as BRAKER3, GALBA and BRAKER2 include costly processing steps for the extrinsic evidence and training of their HMM models.

Tiberius demonstrated the fastest runtime per genome with an average of 1:39h, faster than AUGUSTUS at 2:25h, while Helixer lagged significantly at 8:54h (Table 1, Supplementary Table 4). Tiberius leverages GPU processing across all computational steps of its model, including the HMM. It employs batch parallelization and a parallel implementation of the Viterbi algorithm (see Parallel Viterbi). In contrast, Helixer utilizes the GPU’s computational advantages only for its CNN-LSTM components, lacking GPU integration for its HMM processing. The runtimes of BRAKER3, GALBA, and BRAKER2 were significantly higher, with BRAKER3 being the slowest, with an average runtime of over 2 days.

## Discussion

Tiberius and BRAKER3 both use previously annotated mammalian genomes. BRAKER3 *explicitly* aligns protein sequences against sequences from the target species using several alignment tools. Tiberius, however, only *implicitly* represents the prior knowledge of the annotations *on nucleotide level* in a large number of parameters that are trained with the objective of maximizing a measure of gene prediction accuracy. The fact that Tiberius is even slightly more accurate than BRAKER3, although the latter uses RNA-seq data in addition to a protein database, suggests that the deep learning approach of Tiberius is more effective. Moreover, the new paradigm of representing knowledge within parameters has not yet been subject to the extensive research and refinement seen in traditional pipelines that integrate alignment tools with “shallow” machine learning methods. Consequently, it holds the potential for faster advancements in the future.

The integration of RNA-seq into a genome annotation pipeline has been known to be very beneficial [10] and is even necessary if transcriptome assemblers are used. However, using ten RNA-seq libraries per species has not helped BRAKER3 to surpass the accuracy of Tiberius. We are not aware of a published and benchmarked general-purpose annotation pipeline that is more accurate than BRAKER3 and consequently than Tiberius as an *ab initio* gene finder. When Tiberius could exploit evolutionary information from aligned genomes in *de novo* mode, it even predicted two out of three human genes without error. This approach requires significant bioinformatics resources, as whole genome alignments need to be processed, but is the most accurate among the compared genome annotation methods and does not require any other data besides genomes. Therefore, currently transcriptome sequencing is becoming dispensable if it is only done to increase the average accuracy of mammalian genome annotation. Naturally, in the future, deep learning gene finders that integrate RNA-seq as well may be even more accurate.

### BUSCO as an Accuracy Measure

Although BUSCO completeness is a useful indicator of the sensitivity of predicting the *presence* of genes when annotating a genome, it is not suited to assess gene *structure* accuracy. False positive or missing exons need not affect the BUSCO scores, neither do false positive genes. Furthermore, BUSCO genes are widely conserved, which introduces a bias such that performance weaknesses on less widely conserved genes – the majority – may be undetected. The example of Helixer shows that high BUSCO completeness can go along with a low accuracy on gene level. For example, on human Helixer achieves a very good BUSCO score of about 95%, while only 19% of its predicted genes are correct. In addition, BUSCO scores do not reflect relative performance in the prediction of gene structures: On cow and beluga whale, Helixer achieves a higher BUSCO completeness than BRAKER3 but predicts less than 40% as many genes correctly.

### HMMs for Gene Prediction

From our experiments with deep learning architectures, we draw the conclusion that two things should be customized to the gene prediction task. First, an HMM is a crucial layer to be included in the model architecture because general-purpose sequence-to-sequence models are not competitive without it. Tiberius, Helixer and AUGUSTUS all use an HMM, and AUGUSTUS – now the weakest of the three – still beats all 13 DNA language models tested by Marin *et al*. at the task of human gene prediction. We attribute this to the inductive bias that the HMM introduces by enforcing the common biological knowledge about the admissible label sequences: gene structures obey a regular grammar that is defined by the HMM’s transition graph. Secondly, the choice of a loss function that is adapted to gene prediction is also vital. Whether a base is labeled ‘intron’ or ‘intergenic’ is determined by neighboring exons that may be distant. The precise identification of exon boundaries is therefore particularly important; we achieve this with a custom loss that explicitly penalizes exon boundary errors.

### Limitations of Tiberius and Future Work

It is currently not recommended to use Tiberius to annotate non-vertebrate genomes without retraining it. BRAKER3 has the advantage over Tiberius in that it does not need training gene structures. A practical approach to annotating another clade may be to use BRAKER3 for a selection of genomes with available RNA-seq data, to train Tiberius on the BRAKER3 annotations, and to use Tiberius for the remaining genomes of the clade.

The prediction of alternative splicing and, relatedly, the proper integration of RNA-seq data is not naturally addressed by Tiberius and other sequence-to-sequence models, which find *one* label per position. For this, another approach may have to be found that allows one to model multiple alternative label sequences, each of which obeys the grammar of gene structures with its long-range dependencies.

## Conclusion

*Ab initio* genome annotation can be as accurate as genome annotation based on RNA-seq mapping and protein alignment. When exploiting prior knowledge through homology, a fast, convenient and accurate alternative to aligning individual protein sequences is to represent the prior knowledge in the parameters of a machine learning model instead.

## Acknowledgements

We thank Alisandra Denton for pioneering deep learning in gene finding and for sharing her team’s experience in training Helixer.

## Notes

### Competing Interest Statement

The authors have declared no competing interest.

