## Supplementary Materials for "Tiberius: End-to-End Deep Learning with an HMM for Gene Prediction"

#### 1 Supplementary Figures

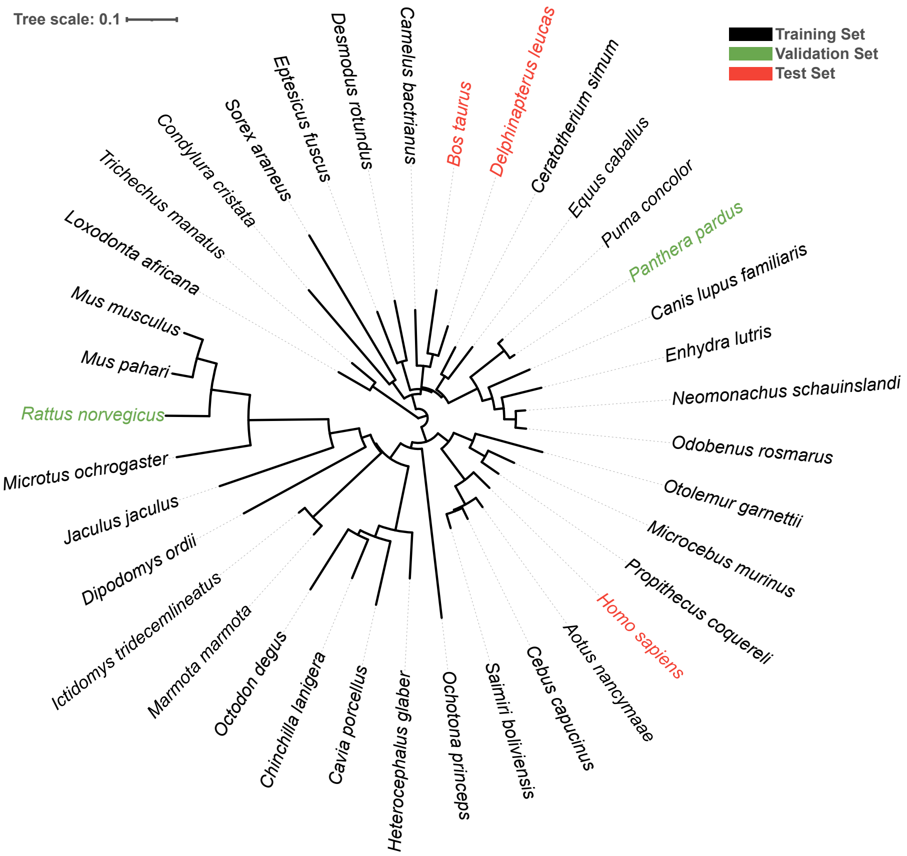


Supplementary Figure S1: Phylogenetic tree derived from UCSC's 241 mammalian Cactus [2] alignments [31] scaled to genomic mutations and illustrating the 37 mammalian species used in the experiments. The Tiberius model was trained on a subset of 32 species (black) and validated on *Rattus norvegicus* and *Panthera pardus* (green), and evaluated on 3 species (red). The tree was visualized with the Interactive Tree Of Life (iTOL) v5 tool [19].

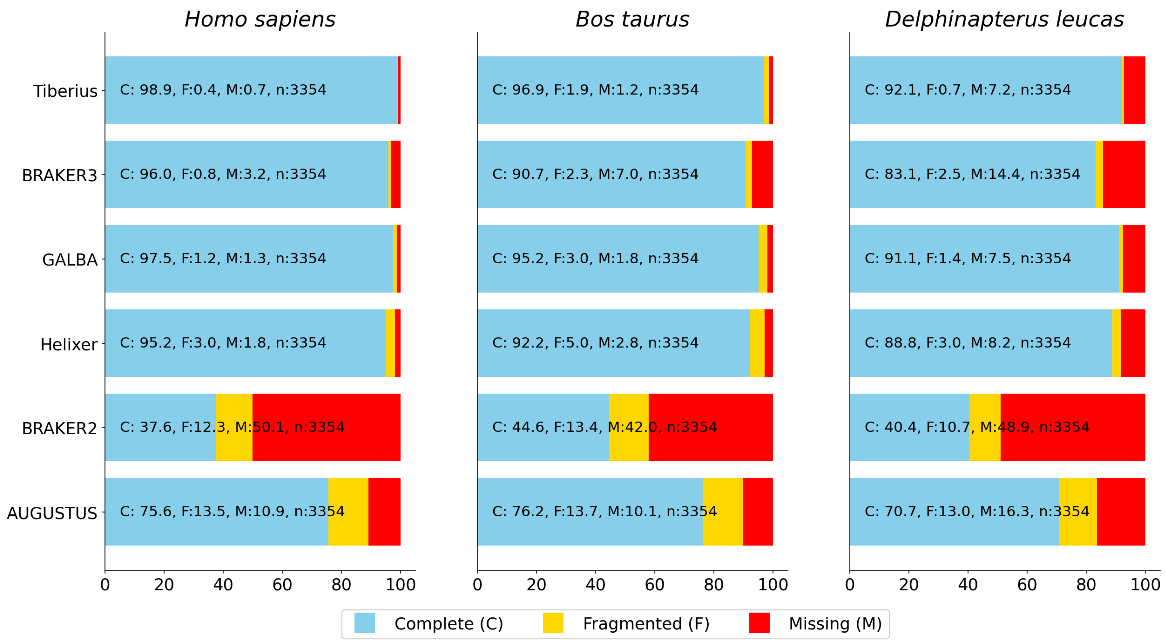


Supplementary Figure S2: BUSCO statistics for test species *Homo sapiens*, *Bos taurus*, and *Delphinapterus leucas*, showing results from six different gene prediction methods. Lineage dataset 'vertebrata_odb10' was used for the analyses.

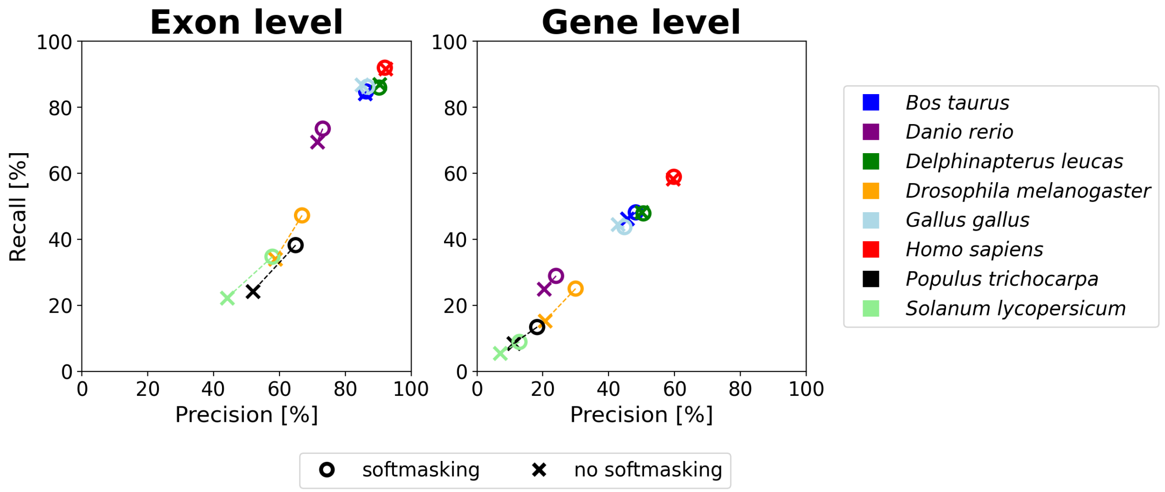


Supplementary Figure S3: Comparison of precision and recall at gene and exon level of the Tiberius model with (o) and without (x) the use of a softmasking track as additional input during training and inference.

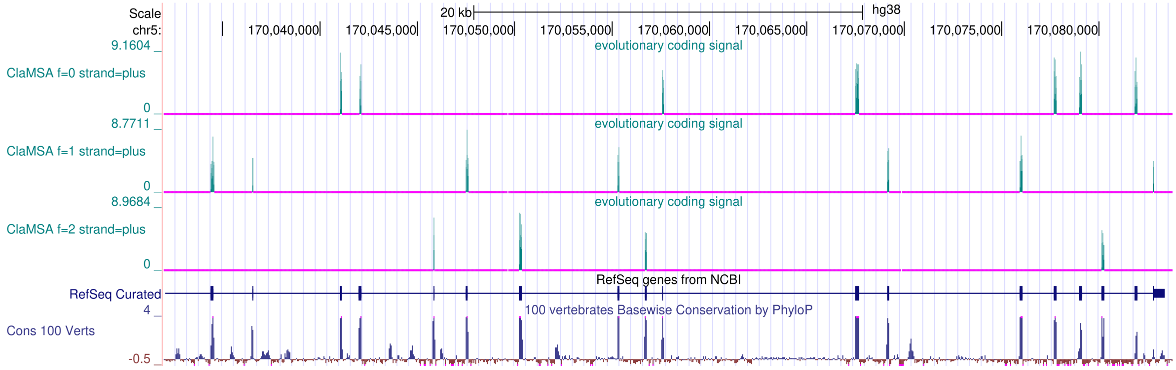


Supplementary Figure S4: ClaMSA positionwise predictions of protein-coding sites in human derived from a 64-way alignment of mammals. Logit values are shown separately for 3 reading frames on the forward strand only. Negative logit values are clipped at 0 (magenta). The plot was created with the UCSC Genome Browser [26]

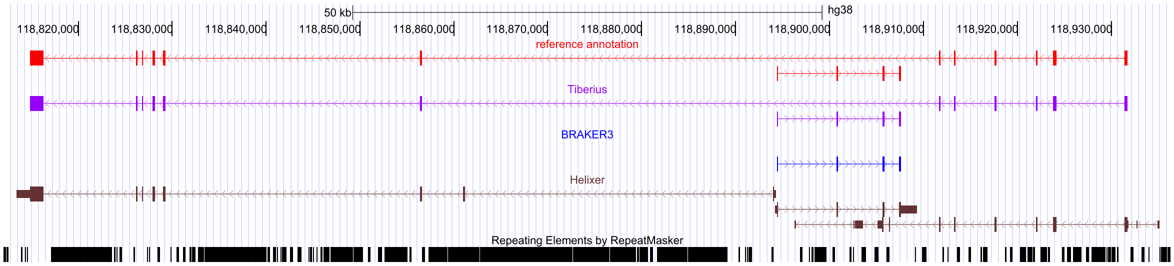


Supplementary Figure S5: Predictions by Tiberius, BRAKER3, and Helixer for a gene locus in the human genome, plotted against a high-quality reference annotation. On the forward strand, all three methods identify the correct gene structure. However, on the reverse strand, only Tiberius accurately predicts the correct gene structure. BRAKER3 fails to detect this gene, while Helixer incorrectly splits it into two shorter genes. The plot was created with the UCSC Genome Browser [26]

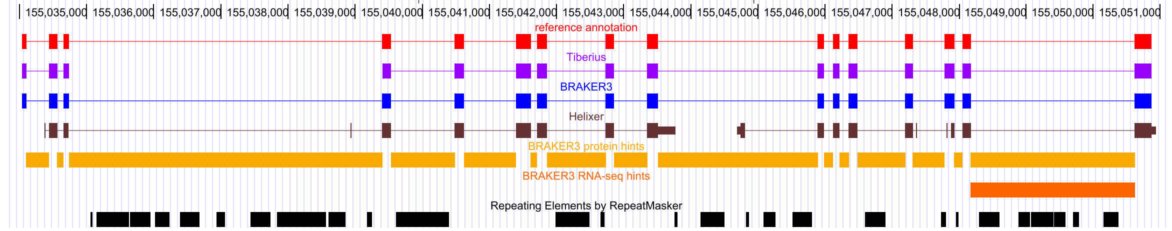


Supplementary Figure S6: Predictions by Tiberius, BRAKER3, and Helixer for a gene locus in the human genome, plotted against a high-quality reference annotation. Tiberius and Helixer ran in *ab initio* mode, while BRAKER3 predicted informed by extrinsic evidence in the form of intron position hints derived from RNA-seq (orange) and protein (yellow) evidence. The plot was created with the UCSC Genome Browser [26]

#### 2 Supplementary Tables

| Species | Accession | Size (Mbp) | #Genes | #Transcripts | #Exons | BUSCO (%) |
|---|---|---|---|---|---|---|
| *Aotus nancymaae* | GCF_000952055.2 | 2,862 | 20,526 | 47,628 | 208,544 | 98.0 |
| *Camelus bactrianus* | GCF_000767855.1 | 1,993 | 19,242 | 28,672 | 195,147 | 96.6 |
| *Canis lupus familiaris* | GCF_000002285.3 | 2,411 | 20,041 | 58,930 | 214,515 | 97.4 |
| *Cavia porcellus* | GCF_000151735.1 | 2,723 | 20,259 | 37,643 | 205,019 | 97.7 |
| *Cebus capucinus* | GCF_001604975.1 | 2,718 | 21,706 | 56,047 | 212,004 | 97.5 |
| *Ceratotherium simum* | GCF_000283155.1 | 2,464 | 20,124 | 33,769 | 200,999 | 99.0 |
| *Chinchilla lanigera* | GCF_000276665.1 | 2,391 | 20,686 | 45,597 | 208,809 | 96.0 |
| *Condylura cristata* | GCF_000260355.1 | 1,770 | 17,965 | 29,249 | 184,170 | 92.1 |
| *Desmodus rotundus* | GCF_002940915.1 | 2,064 | 18,675 | 29,889 | 195,481 | 98.5 |
| *Dipodomys ordii* | GCF_000151885.1 | 2,236 | 19,987 | 29,552 | 194,625 | 94.4 |
| *Enhydra lutris* | GCF_002288905.1 | 2,455 | 19,458 | 36,721 | 205,398 | 99.5 |
| *Eptesicus fuscus* | GCF_000308155.1 | 2,027 | 18,851 | 49,940 | 202,535 | 96.9 |
| *Equus caballus* | GCF_000002305.2 | 2,475 | 20,263 | 36,051 | 204,528 | 95.5 |
| *Heterocephalus glaber* | GCF_000247695.1 | 2,618 | 20,063 | 60,899 | 212,642 | 97.1 |
| *Ictidomys tridecemlineatus* | GCF_000236235.1 | 2,478 | 20,022 | 38,690 | 199,954 | 96.3 |
| *Jaculus jaculus* | GCF_000280705.1 | 2,835 | 18,882 | 25,493 | 191,173 | 96.2 |
| *Loxodonta africana* | GCF_000001905.1 | 3,197 | 21,296 | 41,469 | 203,998 | 97.1 |
| *Marmota marmota* | GCF_001458135.1 | 2,511 | 21,080 | 31,899 | 195,101 | 97.0 |
| *Microcebus murinus* | GCF_000165445.2[1] | 2,487 | 20,671 | 59,172 | 216,624 | 98.3 |
| *Microtus ochrogaster* | GCF_000317375.1 | 2,287 | 20,253 | 37,971 | 201,198 | 98.1 |
| *Mus musculus* | GCF_000001635.26 | 2,819 | 24,228 | 88,511 | 243,350 | 99.7 |
| *Mus pahari* | GCF_900095145.1[2] | 2,475 | 20,649 | 42,393 | 204,649 | 99.0 |
| *Neomonachus schauinslandi* | GCF_002201575.1 | 2,401 | 18,850 | 28,412 | 193,733 | 98.1 |
| *Ochotona princeps* | GCF_000292845.1 | 2,230 | 18,605 | 25,707 | 189,170 | 94.6 |
| *Octodon degus* | GCF_000260255.1 | 2,996 | 20,774 | 42,587 | 203,072 | 96.7 |
| *Odobenus rosmarus* | GCF_000321225.1 | 2,400 | 19,472 | 31,430 | 201,082 | 98.6 |
| *Otolemur garnettii* | GCF_000181295.1 | 2,520 | 19,666 | 32,664 | 195,646 | 97.5 |
| *Panthera pardus* | GCF_001857705.1[3] | 2,578 | 20,422 | 58,058 | 216,459 | 99.2 |
| *Propithecus coquereli* | GCF_000956105.1 | 2,798 | 19,484 | 28,287 | 189,571 | 92.5 |
| *Puma concolor* | GCF_003327715.1 | 2,433 | 17,707 | 23,406 | 182,810 | 92.6 |
| *Rattus norvegicus* | GCF_000001895.5 | 2,870 | 23,409 | 56,583 | 226,229 | 97.8 |
| *Saimiri boliviensis* | GCF_000235385.1 | 2,609 | 19,833 | 36,325 | 201,157 | 96.8 |
| *Sorex araneus* | GCF_000181275.1 | 2,423 | 19,398 | 23,609 | 189,074 | 96.1 |
| *Trichechus manatus* | GCF_000243295.1 | 3,104 | 19,180 | 36,478 | 199,593 | 96.8 |

**Supplementary Table S1.** Summary of genomic characteristics for the dataset, consisting of 34 mammalian species, that were used for **training** and **validating** the Tiberius model. The table shows genome size in Megabases (Mbp), the number of genes, transcripts, protein-coding exons, and the BUSCO completeness score of the reference annotation. Lineage dataset 'vertebrata_odb10' was used for the BUSCO analyses. In all cases, if both a GenBank and a RefSeq genome annotation existed, we chose to use the RefSeq annotation. For three species, this results in a difference between accession numbers for annotation and genome assembly. The deviating genome assembly accessions are: [1]) GCA_000165445.3, [2]) GCA_900095145.2, [3]) GCA_001857705.1.

| Species | Accession | Size (Mbp) | #Genes | #Transcripts | #Exons | BUSCO (%) |
|---|---|---|---|---|---|---|
| *Homo sapiens* | GCF_000001405.40 | 3,257 | 19,411 | 67,043 | 215,455 | 100.0 |
| *Bos taurus* | GCF_000003205.7 | 2,725 | 21,427 | 28,999 | 200,657 | 96.1 |
| *Delphinapterus leucas* | GCF_002288925.1 | 2,359 | 18,524 | 49,701 | 207,096 | 98.4 |

**Supplementary Table S2.** Summary of genomic characteristics for the dataset, consisting of 3 mammalian species, that were used for **testing** the Tiberius model. The table showes genome size in Megabases (Mbp), the number of genes, transcripts, protein-coding exons, and the BUSCO completeness score of the reference annotation. Lineage dataset 'vertebrata_odb10' was used for the BUSCO analyses.

| Species | SRA ID | #spots | #bases |
|---|---|---|---|
| *Homo sapiens* | DRR544106 | 56,983,810 | 17,209,110,620 |
| | SRR22425744 | 24,124,166 | 7,237,249,800 |
| | SRR28771423 | 33,333,805 | 10,066,809,110 |
| | SRR28784504 | 59,935,845 | 17,980,753,500 |
| | SRR28789035 | 42,367,420 | 12,794,960,840 |
| | SRR28805051 | 38,393,690 | 11,518,107,000 |
| | SRR28817629 | 63,195,932 | 12,765,578,264 |
| | SRR28876548 | 69,731,391 | 14,058,415,314 |
| | SRR28891528 | 37,131,776 | 11,139,532,800 |
| | SRR28920169 | 12,197,090 | 3,659,127,000 |
| *Bos taurus* | SRR26343080 | 36,208,362 | 10,862,508,600 |
| | SRR26343251 | 38,912,939 | 11,673,881,700 |
| | SRR26343311 | 33,544,979 | 10,063,493,700 |
| | SRR26343718 | 40,970,110 | 12,291,033,000 |
| | SRR26343784 | 33,604,220 | 10,081,266,000 |
| | SRR27959844 | 36,134,498 | 10,840,349,400 |
| | SRR27959903 | 35,640,738 | 10,692,221,400 |
| | SRR28355215 | 47,218,711 | 14,260,050,722 |
| | SRR28762329 | 36,672,650 | 11,075,140,300 |
| | SRR28762335 | 39,626,087 | 11,967,078,274 |
| *Delphinapterus leucas* | SRR5282283 | 30,325,796 | 4,548,869,400 |
| | SRR5282284 | 33,217,830 | 4,982,674,500 |
| | SRR5282288 | 29,404,272 | 4,410,640,800 |
| | SRR5282294 | 31,046,128 | 4,656,919,200 |
| | SRR5990718 | 11,916,084 | 4,067,286,534 |
| | SRR6181296 | 6,312,007 | 951,737,487 |
| | SRR6181297 | 6,184,859 | 932,537,966 |
| | SRR6181300 | 5,707,619 | 858,497,117 |
| | SRR6181301 | 5,845,122 | 879,167,100 |
| | SRR6181303 | 6,148,542 | 924,655,926 |

**Supplementary Table S3.** List of all RNA-seq libraries used for the test species in the experiments of BRAKER3. It includes the ID of the library from the Sequence Read Archive, the number of spots, and the number of bases.

| | Runtime (h:m) | | | |
|---|---|---|---|---|
| | *H. sapiens* | *B. taurus* | *D. leucas* | Average |
| Tiberius | **1:56** | **1:43** | **1:18** | **1:39** |
| Helixer | 9:28 | 8:56 | 8:18 | 8:54 |
| AUGUSTUS | 2:49 | 2:24 | 2:01 | 2:25 |
| BRAKER3 | 62:30 | 43:39 | 40:30 | 48:53 |
| GALBA | 35:52 | 35:53 | 33:51 | 35:12 |
| BRAKER2 | 16:36 | 15:54 | 14:19 | 15:36 |

**Supplementary Table S4.** Runtime comparison for the inference of gene structures using Tiberius, BRAKER3, GALBA, Helixer, BRAKER2, and AUGUSTUS. BRAKER3, GALBA, BRAKER2, and AUGUSTUS were executed on a CPU utilizing 48 parallel threads, while Helixer and Tiberius were run on a GPU node with one A100 GPU.

| | | Runtime (h:m) | | | |
|---|---|---|---|---|---|
| Hardware | HMM parallelization | *H. sapiens* | *B. taurus* | *D. leucas* | Average |
| 96-core CPU | Yes | 28:08 | 25:15 | 20:10 | 24:31 |
| A100 GPU | No | 34:28 | 31:53 | 19:36 | 28:39 |
| A100 GPU | Yes | **1:56** | **1:43** | **1:18** | **1:39** |

**Supplementary Table S5.** Whole genome runtime comparison for the inference of gene structures using Tiberius running on different hardware (A100 GPU vs 96 core CPU) and with and without using parallelization of the HMM Layer.

| | Exon | | | Gene | | |
|---|---|---|---|---|---|---|
| | Recall | Precision | F1 | Recall | Precision | F1 |
| *Homo sapiens* | | | | | | |
| Tiberius | **92.81** | 92.63 | **92.72** | 63.08 | 60.79 | 61.91 |
| Tiberius *de novo* | 92.00 | **93.29** | 92.64 | **66.55** | **64.47** | **65.49** |
| BRAKER3 | 82.80 | 92.25 | 87.27 | 61.29 | 55.37 | 58.18 |
| GALBA | 89.62 | 87.61 | 88.60 | 50.68 | 41.52 | 45.64 |
| Helixer | 85.06 | 62.02 | 71.74 | 22.57 | 19.11 | 20.70 |
| BRAKER2 | 41.63 | 58.25 | 48.56 | 22.95 | 12.73 | 16.38 |
| AUGUSTUS | 69.82 | 69.28 | 69.55 | 14.44 | 13.29 | 13.84 |
| *Bos taurus* | | | | | | |
| Tiberius | **86.13** | 87.46 | **86.79** | **50.42** | 51.35 | **50.88** |
| BRAKER3 | 71.82 | **90.50** | 80.09 | 46.50 | **55.16** | 50.46 |
| GALBA | 82.30 | 83.93 | 83.11 | 40.11 | 37.46 | 38.74 |
| Helixer | 78.43 | 66.01 | 71.69 | 17.96 | 19.17 | 18.55 |
| BRAKER2 | 44.32 | 53.25 | 48.38 | 19.82 | 10.84 | 14.01 |
| AUGUSTUS | 65.70 | 66.09 | 65.89 | 11.44 | 12.21 | 11.81 |
| *Delphinapterus leucas* | | | | | | |
| Tiberius | **88.04** | 91.34 | **89.66** | **51.28** | 53.59 | 52.41 |
| BRAKER3 | 73.85 | **92.45** | 82.11 | 50.79 | **54.37** | **52.52** |
| GALBA | 86.21 | 87.51 | 86.86 | 43.04 | 39.32 | 41.10 |
| Helixer | 81.04 | 70.38 | 75.33 | 18.62 | 18.94 | 18.78 |
| BRAKER2 | 43.42 | 57.00 | 49.29 | 20.35 | 10.94 | 14.23 |
| AUGUSTUS | 67.21 | 65.90 | 66.55 | 12.00 | 11.10 | 11.53 |
| Average | | | | | | |
| Tiberius | **88.99** | 90.48 | **89.72** | **54.93** | **55.24** | **55.07** |
| BRAKER3 | 76.16 | **91.73** | 83.15 | 52.86 | 54.97 | 53.72 |
| GALBA | 86.04 | 86.35 | 86.19 | 44.61 | 39.43 | 41.83 |
| Helixer | 81.51 | 66.14 | 72.92 | 19.72 | 19.07 | 19.34 |
| BRAKER2 | 43.12 | 56.17 | 48.74 | 21.04 | 11.50 | 14.87 |
| AUGUSTUS | 67.58 | 67.09 | 67.33 | 12.63 | 12.20 | 12.40 |

**Supplementary Table S6.** Gene and exon level precision and recall of predictions by Tiberius, BRAKER3, GALBA, Helixer, BRAKER2, and AUGUSTUS. Tiberius, Helixer, and AUGUSTUS are *ab initio* methods, while GALBA used protein sequences from a three related species, BRAKER2 used a large protein database, and BRAKER3 used a large protein database with RNA-seq data. Additionally, Tiberius was run in *de novo* mode for the human genome, using evolutionary evidence that was generated with ClaMSA.

| | Exon | | | Gene | | |
|---|---|---|---|---|---|---|
| | Recall | Precision | F1 | Recall | Precision | F1 |
| *Homo sapiens* | | | | | | |
| Tiberius | 92.81 | 92.63 | 92.72 | 63.08 | 60.79 | 61.91 |
| Tiberius_preHMM | 91.50 | 92.35 | 91.92 | 60.08 | 58.68 | 59.37 |
| Tiberius_no_sm | 92.09 | 91.63 | 91.86 | 59.90 | 56.89 | 58.36 |
| Tiberius_small | 90.02 | 90.85 | 90.43 | 55.35 | 51.24 | 53.22 |
| Tiberius_CCE | 89.08 | 90.20 | 89.64 | 48.86 | 49.04 | 48.95 |
| Tiberius_5class | 86.37 | 86.47 | 86.42 | 46.54 | 44.48 | 45.49 |
| Tiberius_5class_CCE | 80.38 | 81.83 | 81.10 | 34.00 | 34.76 | 34.38 |
| *Bos taurus* | | | | | | |
| Tiberius | 86.13 | 87.46 | 86.79 | 50.42 | 51.35 | 50.88 |
| Tiberius_preHMM | 84.73 | 87.04 | 85.87 | 47.64 | 49.44 | 48.52 |
| Tiberius_no_sm | 84.57 | 85.61 | 85.09 | 46.55 | 45.53 | 46.03 |
| Tiberius_small | 83.34 | 85.81 | 84.56 | 44.48 | 44.55 | 44.51 |
| Tiberius_CCE | 82.55 | 85.49 | 83.99 | 40.01 | 43.04 | 41.47 |
| Tiberius_5class | 79.60 | 81.33 | 80.46 | 38.75 | 39.15 | 38.95 |
| Tiberius_5class_CCE | 74.61 | 77.76 | 76.15 | 29.79 | 32.43 | 31.05 |
| *Delphinapterus leucas* | | | | | | |
| Tiberius | 88.04 | 91.34 | 89.66 | 51.28 | 53.59 | 52.41 |
| Tiberius_preHMM | 85.59 | 90.85 | 88.14 | 47.75 | 51.54 | 49.57 |
| Tiberius_no_sm | 85.56 | 90.17 | 87.80 | 47.32 | 49.96 | 48.60 |
| Tiberius_small | 84.83 | 89.65 | 87.17 | 44.22 | 45.78 | 44.99 |
| Tiberius_CCE | 83.51 | 88.59 | 85.98 | 39.03 | 42.36 | 40.63 |
| Tiberius_5class | 80.26 | 85.23 | 82.67 | 36.48 | 38.98 | 37.69 |
| Tiberius_5class_CCE | 74.97 | 80.65 | 77.71 | 27.36 | 30.54 | 28.86 |
| Average | | | | | | |
| Tiberius | 88.99 | 90.48 | 89.72 | 54.93 | 55.24 | 55.07 |
| Tiberius_preHMM | 87.27 | 90.08 | 88.64 | 51.82 | 53.22 | 52.49 |
| Tiberius_no_sm | 87.41 | 89.14 | 88.25 | 51.26 | 50.79 | 51.00 |
| Tiberius_small | 86.06 | 88.77 | 87.39 | 48.02 | 47.19 | 47.57 |
| Tiberius_CCE | 85.05 | 88.09 | 86.54 | 42.63 | 44.81 | 43.68 |
| Tiberius_5class | 82.08 | 84.34 | 83.18 | 40.59 | 40.87 | 40.71 |
| Tiberius_5class_CCE | 76.65 | 80.08 | 78.32 | 30.38 | 32.58 | 31.43 |

**Supplementary Table S7.** Gene and exon level precision and recall of predictions of ablation studies from Tiberius, where the pre-HMM model of Tiberius was trained from scratch with key features of Tiberius changed.

| Tile length | Exon | | | Gene | | |
|---|---|---|---|---|---|---|
| | Recall | Precision | F1 | Recall | Precision | F1 |
| *Panthera pardus* | | | | | | |
| 9999 | 34.51 | 65.48 | 45.20 | 26.05 | 28.14 | 27.05 |
| 29,997 | 56.94 | 75.66 | 64.98 | 35.10 | 36.05 | 35.57 |
| 99,999 | 77.04 | 83.08 | 79.95 | 39.51 | 41.25 | 40.36 |
| 199,998 | 82.33 | 85.31 | 83.79 | 39.69 | 42.43 | 41.01 |
| 500,004 | 83.54 | 86.45 | 84.97 | 38.66 | 42.70 | 40.58 |
| 1,000,008 | 80.61 | 86.60 | 83.50 | 37.07 | 42.87 | 39.76 |
| *Rattus norvegicus* | | | | | | |
| 9999 | 37.68 | 68.73 | 48.67 | 33.68 | 35.05 | 34.35 |
| 29,997 | 60.36 | 78.48 | 68.24 | 43.04 | 42.61 | 42.82 |
| 99,999 | 78.59 | 84.32 | 81.35 | 46.57 | 46.23 | 46.40 |
| 199,998 | 83.05 | 85.73 | 84.37 | 46.73 | 47.00 | 46.86 |
| 500,004 | 84.62 | 86.45 | 85.53 | 46.75 | 47.46 | 47.10 |
| 1,000,008 | 84.45 | 86.55 | 85.49 | 46.44 | 47.51 | 46.97 |
| Average | | | | | | |
| 9999 | 36.09 | 67.11 | 46.94 | 29.87 | 31.59 | 30.70 |
| 29,997 | 58.65 | 77.07 | 66.61 | 39.07 | 39.33 | 39.20 |
| 99,999 | 77.81 | 83.70 | 80.65 | 43.04 | 43.74 | 43.38 |
| 199,998 | 82.69 | 85.52 | 84.08 | 43.21 | 44.72 | 43.94 |
| 500,004 | 84.08 | 86.45 | 85.25 | 42.70 | 45.08 | 43.84 |
| 1,000,008 | 82.53 | 86.57 | 84.49 | 41.75 | 45.19 | 43.36 |

**Supplementary Table S8.** Gene and exon level precision and recall of predictions by Tiberius for the validation species using different tile lengths during inference. The model training and weights remained the same, only the tile lengths during inference differed in these experiments.

### 3 Supplementary Methods

#### 3.1 Assembly Datasets

The (full) genome assemblies were extracted from the Zoonomia genome alignment of mammals [1]. RepeatModeler2 [9] was used to generate species-specific repeat libraries. RepeatMasker (`http://www.repeatmasker.org`) was then used to softmask repeats, followed by Tandem Repeats Finder [4] to ensure more thorough masking. Reference annotations were retrieved from the NCBI in GFF3 format and were split were split into a GTF file with protein-coding gene annotations, and into a file with pseudogenes in GFF3 format.

##### 3.1.1 Genome Extraction

Genomes were extracted from the Zoonomia alignment with hal2fasta v2.1 as follows:

```
cat species.names | \
while read -r species
do
  echo "$species"
  if [ ! -f ${species}.fa.gz ]; then
    hal2fasta mammals.hal $species | gzip -c > ${species}.fa.gz
  fi
done
```

##### 3.1.2 Repeat Masking

RepeatModeler2 version v2.0.2 was used to construct a species-specific database (here: `$DB`) from each species genome in FASTA format (here `$GENOME`):

```
BuildDatabase -name ${DB} ${GENOME}
RepeatModeler -database ${DB} -pa 72 -LTRStruct
```

Subsequently, RepeatMasker v4.1.4 was used with that repeat library to initially mask the genome, resulting in a masked FASTA file `$SPECIES.fa.masked`:

```
RepeatMasker -pa 72 -lib ${DB} -xsmall ${SPECIES}.fa
```

Since that masking has left several repetitive regions unmasked, an additional masking step with Tandem Repeats Finder was applied, resulting in a file `$SPECIES.fa.combined.masked` (data parallelization was applied):

```
mkdir trf
cd trf
ln -s ../genome.fa.masked genome.fa
splitMfasta.pl --minsize=25000000 genome.fa
ls genome.split.*.fa | parallel 'trf {} 2 7 7 80 10 50 500 -d -m -h &> {}.log'
ls genome.split.*.fa.2.7.7.80.10.50.500.dat | parallel 'parseTrfOutput.py {} --minCopies 1 \
    --statistics {}.STATS > {}.raw.gff 2> {}.parsedLog'
ls genome.split.*.fa.2.7.7.80.10.50.500.dat.raw.gff | parallel 'sort -k1,1 -k4,4n -k5,5n {} \
    > {}.sorted 2> {}.sortLog'
FILES=genome.split.*.fa.2.7.7.80.10.50.500.dat.raw.gff.sorted
for f in $FILES do
    bedtools merge -i $f | awk 'BEGIN{OFS="\t"} {print $1,"trf","repeat",$2+1,$3,".",".",".","."}' \
        > $f.merged.gff 2> $f.bedtools_merge.log
done
ls genome.split.*.fa | parallel 'bedtools maskfasta -fi {} -bed \
    {}.2.7.7.80.10.50.500.dat.raw.gff.sorted.merged.gff -fo {}.combined.masked -soft &> {}.bedools_mask.log'
cat genome.split.*.fa.combined.masked > genome.fa.combined.masked
```

The software versions used: trf 4.0.9, splitMfasta.pl from AUGUSTUS v3.5.0, bedtools v2.30.0.

##### 3.1.3 Reference Annotations

Reference annotations were retrieved from the NCBI (accession numbers in Tables 1 and 2). These reference annotations were in the GFF3 format and were post-processed according to the protocol used to format reference annotations in the BRAKER2 publication [5], documented at `https://github.com/gatech-genemark/EukSpecies-BRAKER2` (required scripts used in the command lines shown below are also available there).

For the annotation of *Homo sapiens*, in rarely occurring hash marks were deleted from the GFF3 lines of gene features. Also, for *Mus musculus* and *Rattus norvegicus*, hash marks and percentage characters were removed. These steps did not affect gene structures. In short, the commands (for all species) were as follows.

```
zcat genome.fa  | grep '^>' > defline
zcat ${ANNOT}.gff.gz | perl -pe 's/\%/_/g;' | perl -ne 'if(not(m/^\#/)){s/\#/_/g;} print $_;' \
```

```
    > ${ANNOT}.f.gff
cat defline | cut -f1 -d ' ' | cut -b2- > z
paste z z > list.tbl
echo "##gff-version 3" > annot.gff3
probuild --stat_fasta --seq genome.fasta | cut -f1,2 | tr -d '>' |  grep -v '^$' | \
awk '{print "##sequence-region  " $1 "  1 " $2}\
' >> annot.gff3
cat ${ANNOT}.f.gff | grep -v "#" >> annot.gff3
gt  gff3  -force  -tidy  -sort  -retainids  -checkids  -o tmp_annot.gff3  annot.gff3
mv tmp_annot.gff3 annot.gff3
select_pseudo_from_nice_gff3.pl annot.gff3 pseudo.gff3
enrich_gff.pl --in annot.gff3 --out tmp_annot.gff3 --cds --seq genome.fasta --v --warnings
mv tmp_annot.gff3 annot.gff3
gff3_to_gtf.pl annot.gff3 annot.gtf
```

##### 3.2 Running Tiberius

Tiberius was run using a softmasked genome `genome.fa` and the model weights `tiberius_weights` from `https://bioinf.uni-greifswald.de/bioinf/tiberius/models/tiberius_weights.tgz`:

```
tiberius.py --genome genome.fa --model default_model_weights \
    --out tiberius.gtf
```

Tiberius in *de novo* mode was run using a softmasked genome, ClaMSA data generated as documented in Supplementary Methods 3.7, and the model weights `tiberius_denovo_weights` from `https://bioinf.uni-greifswald.de/bioinf/tiberius/models/tiberius_denovo_weights.tgz`:

```
tiberius.py --genome genome.fa --model tiberius_denovo_weights \
    --clamsa path/to/clamsa/data  --out tiberius.gtf
```

##### 3.3 Running Helixer

Helixer v0.3.3 was executed within a Singularity [17] container obtained from `docker://gglyptodon/helixer-docker:helixer_v0.3.3_cuda_11.8.0-cudnn8`. The tool was run on all three test species using the recommended parameters and model weights for vertebrates on a machine with a single A100 (80Gb) GPU.

```
 Helixer.py --lineage vertebrate --fasta-path genome.fa --gff-output-path helixer.gff \
       --subsequence-length 213840 --overlap-offset 106920 --overlap-core-length 160380
```

##### 3.4 Running BRAKER

BRAKER v3.0.8 was installed from GitHub (`https://github.com/Gaius-Augustus/BRAKER`) and for all three test species run as:
BRAKER2:

```
braker.pl --genome=genome.softmasked.fasta --prot_seq=proteins.fa --threads=72
```

BRAKER3:

```
braker.pl --genome=genome.softmasked.fasta --prot_seq=proteins.fa --rnaseq_sets_ids=RNA_Seq_IDS \
    --threads=72
```

The RNA-seq data was automatically downloaded and prepared with BRAKER. The protein database (`proteins.fa`) used was the vertebrate partition of OrthoDB with exclusion of *Hominidae* (*Homo sapiens*), *Ruminantia* (*Bos taurus*), or *Cetacea* (*Delphinapterus leucas*), depending on the target species. The protein databases were prepared with orthodb-clades (`https://github.com/tomasbruna/orthodb-clades`). For example, the following command was used:

```
selectClade.py  orthodb-clades/clades/Vertebrata.fa orthodb-clades/orthodb/levels.tab \
       orthodb-clades/orthodb/level2species.tab Vertebrata --exclude Hominidae > proteins.fa
```

##### 3.5 Running GALBA

GALBA v1.0.11 was executed within a Singularity container obtained from `docker://katharinahoff/galba-notebook:latest` and run as:

```
galba.pl  --genome=genome.softmasked.fasta --prot_seq=proteins.fa --threads 72
```

For each test species, we selected the protein sequences from the annotations of three closely related species from the set of training genomes for Tiberius (Table 1). For *Bos taurus* and *Delphinapterus leucas*, we selected the proteins from *Camelus bactrianus*, *Ceratotherium simum*, and *Odobenus rosmarus*. For *Homo sapiens*, we selected *Microcebus murinus*, *Otolemur garnettii*, *Saimiri boliviensis*.

#### 3.6 BUSCO

BUSCO v5.4.4 scores were computed from the translated protein sequences `proteins.fa` for each gene prediction:

```
busco -m protein -i proteins.fa -o busco -l vertebrata -f -c 48
```

**Accuracy evaluation**
The performance measurements were computed using scripts from the BRAKER scripts:

```
compute_accuracies.sh ref_annot.gtf pseudo.gff3 gene_set.gtf gene cds
```

#### 3.7 Evolutionary Evidence

As an extension to the evidence from the target genome alone (*ab initio* setting) we tested, how *evolutionary evidence* from the comparison of multiple unannoated genomes can further help to increase accuracy. This setting in which multiple genomes are used for gene prediction has been termed *de novo* gene prediction [11]. Note that this problem statement requires neither RNA-Seq data nor a database of protein sequences.

**ClaMSA**
The ClaMSA tool uses gradient-based machine learning to discriminatively train classical models of molecular evolution. It was originally designed to classify entire nucleotide MSAs into coding and noncoding [24].

**Extension and training.** We here describe how ClaMSA was trained and used in this study. Note that a retraining of ClaMSA for other clades is likely not necessary. However, we have not tested how the genome alignment program (here Progressive Cactus) is influencing the results. The functionality of ClaMSA was extended for the individual classification of each alignment column. In addition, an option to input MAF formatted multiple alignments to ClaMSA was implemented. ClaMSA was then trained on the alignment of 41 primate genomes extracted from the alignment generated in [1]. This relatively lightweight model consists of 4 time-reversible $64 \times 64$ codon rate matrices and a small multi-layer perceptron with 2 layers that have 16 hidden units each, totaling to 8,702 trainable parameters. Because the number of parameters of ClaMSA is small and as its parameters were previously shown to be applicable cross-clades (vertebrates parameters work well on flies and vice versa) [24] we chose to train ClaMSA only on a small set of genes. As training labels, we used NCBI's RefSeq annotation of human chromosome 17 with a total of 291,379 positive (coding) sites and a ratio of 6.1 negatives per positive. After training, which took about 8 hours, the test accuracy of the classification of individual sites as coding or noncoding was 93.5%.

**ClaMSA sitewise predictions.** We selected 64 species ("assorted 64") that are a maximally diverse superset of the 37 species displayed in 1 and a subset of the 240 eutherian mammalian species of [1] (Supplementary files assorted64.txt, assorted64.pdf). The HAL formatted alignment of the Zoonomia project [1] was converted with `cactus-hal2maf` [2] into MAF formatted files, and restricted to the 64 assorted species, thus using the human genome as reference (see Supplementary files). ClaMSA was then run in prediction mode to classify all aligned sites in all 64 aligned genomes. It thereby considers a single triplet of aligned nucleotides at a time and outputs a probability $p$ that the site is coding, making the simplifying assumption that a site is either coding in all genomes or in none. This yields WIG (wiggle) formatted files that contain for each genome and strand a value logit$(p)$. Supplementary Figure 4 shows a genome browser screenshot with the ClaMSA tracks. The essential command lines are shown below.

**Tiberius using ClaMSA**
To prepare the ClaMSA output for Tiberius, we summarize the sitewise predictions into four values per position (two per strand). The first value for each position and strand is the mean of logit probabilities for codon start positions from the ClaMSA output. If no ClaMSA information is available, we use $\ln(-5)$ as a lower bound. The second value is the number of probabilities available for this position. We integrate these four values into the Tiberius model by stacking them to the input of each position, maintaining the rest of the model architecture (Figure 1). We trained this modified model from scratch using the same procedure described in Tiberius Training but we reduced the learning rate to $10^{-5}$ during fine-tuning.

##### 3.7.1 Training

The training of ClaMSA requires alignment files in binary TensorFlow records (TFRecords) format, where the alignment columns are annotated with regard to whether they are coding. Such files were created as follows. First, the HAL formatted alignment of 240 eutherian species was projected to the subclade of 41 primates and converted to MAF format with a command that is analogous to the `cactus-hal2maf` described below for predictions. Then, only the part of the alignment that includes human chromosome 17 was annotated with converted to FASTA with the following command using a script included with ClaMSA:

```
coding-labeled-MSAs.py --inmsa chr17-primates.maf.gz --refspecies Homo_sapiens  \
    --anno hg38.chr17.refseq.CDS.gtf  --outmsa chr17-primates.fa.gz
```

To compile a training set in TFRecord file format the following command was used to convert the annotated FASTA file to TFRecods:

```
clamsa.py convert fasta chr17-primates.fa.gz --sitewise --tf_out_dir . --refid 0 \
      --splits '{"train": -1, "test": 3000, "val": 2000}' --use_codons \
      --basename primates_human_chr17 --clades primates.nwk --no_codon_alignment
```

Then ClaMSA was trained on the TFRecord data set generated by the previous command (a version of ClaMSA functionally equivalent to commit 95d8d6c on `https://github.com/Gaius-Augustus/clamsa`).

```
clamsa.py train --clades diverse64.nwk --use_codons --sitewise --classify --basenames diverse64_human_chr17 \
    --split_specification '{
```

```
        "train": {"name": "train", "wanted_models": [0], "interweave_models": true, "repeat_models": [true, true]},
        "val"  : {"name": "val",   "wanted_models": [0], "interweave_models": true, "repeat_models": [false, false]},
        "test" : {"name": "test",  "wanted_models": [0], "interweave_models": true, "repeat_models": [false, false]}
  }' \
  --model_hyperparameters '{
        "tcmc_dNdS_class" : {
            "sparse_rates" : [false],
            "tcmc_models": [4],
            "dense_dimension": [16]
        }
    }' \
  --batch_size 128 --batches_per_epoch 100 --epochs 150 --saved_weights_basedir saved_weights
```

**3.7.2 Sitewise Prediction**

The following format was used to project the multiple genome alignment to assorted 64 genomes referenced by each human chromosome and to convert it to MAF format.

```
cactus-hal2maf js --workDir wd 241-mammalian-2020v2.hal assorted64-${CHR}.maf.gz --refGenome Homo_sapiens \
  --chunkSize 100000 --noAncestors --refSequence $CHR --filterGapCausingDupes --dupeMode single --batchCores 2
  --targetGenomes Aotus_nancymaae,Camelus_bactrianus,Canis_lupus_familiaris,Cavia_porcellus,Cebus_capucinus,\
  Ceratotherium_simum,Chinchilla_lanigera,Condylura_cristata,Desmodus_rotundus,Dipodomys_ordii,Enhydra_lutris,\
  Eptesicus_fuscus,Equus_caballus,Heterocephalus_glaber,Ictidomys_tridecemlineatus,Jaculus_jaculus,\
  Loxodonta_africana,Marmota_marmota,Microcebus_murinus,Microtus_ochrogaster,Mus_musculus,Mus_pahari,\
  Neomonachus_schauinslandi,Ochotona_princeps,Octodon_degus,Odobenus_rosmarus,Otolemur_garnettii,Panthera_pardus,\
  Propithecus_coquereli,Puma_concolor,Rattus_norvegicus,Saimiri_boliviensis,Sorex_araneus,\
  Trichechus_manatus,Bos_taurus,Delphinapterus_leucas,Homo_sapiens,Microgale_talazaci,Erinaceus_europaeus,\
  Elephantulus_edwardii,Ctenodactylus_gundi,Tupaia_chinensis,Tamandua_tetradactyla,Muscardinus_avellanarius,\
  Solenodon_paradoxus,Nannospalax_galili,Petromus_typicus,Crocidura_indochinensis,Chrysochloris_asiatica,\
  Heterohyrax_brucei,Craseonycteris_thonglongyai,Castor_canadensis,Manis_javanica,Uropsilus_gracilis,\
  Catagonus_wagneri,Acomys_cahirinus,Aplodontia_rufa,Dasypus_novemcinctus,Rousettus_aegyptiacus,\
  Sigmodon_hispidus,Zapus_hudsonius,Galeopterus_variegatus,Lepus_americanus,Orycteropus_afer
```

The following command generates WIG formatted files with ClaMSA predictions (logit values) for all 64 genomes on either strand and in 3 phases. The input MAF files were split into chunks for data parallelization.

```
clamsa.py predict maf $CHR/{CHUNK}.maf.gz --clades assorted64.nwk --sitewise --use_codons \
        --model_ids '{ "diverse64mammals" : "mammals-sitewise" }' --output_all_species --logits \
        --out $OUTD/clamsa.$CHR.$CHUNK.
```

#### 3.8 F1-Loss

The estimated F1-score can be computed for an output class $i$ by using the expected values of precision and recall, similarly described by Yacouby and Axman [32]. Let $Y \in \{0,1\}^{(T,K)}$ be the matrix of true labels and $\hat{Y} \in [0,1]^{(T,K)}$ be the predictions. We define the expected predicted positives as $\text{EPP}_i = \sum_{j=1}^{T} \hat{Y}[j,i]$, the actual positives as $\text{AP}_i = \sum_{j=1}^{T} Y[j,i]$, and the expected true positives as $\text{ETP}_i = \sum_{j=1}^{T}(Y[j,i] \cdot \hat{Y}[j,i])$. The estimated precision $\widehat{\text{Pr}}$ and the estimated recall $\widehat{\text{Rc}}$ are then calculated for the class as $\widehat{\text{Pr}}_i = \frac{\text{ETP}_i}{\text{EPP}_i}$ and $\widehat{\text{Rc}}_i = \frac{\text{ETP}_i}{\text{AP}_i}$. The estimated F1-score is then derived as the harmonic mean of the estimated precision and recall:

$$\widehat{\text{F1}}_i = 2 \cdot \frac{\widehat{\text{Pr}}_i \cdot \widehat{\text{Rc}}_i}{\widehat{\text{Pr}}_i + \widehat{\text{Rc}}_i}.$$

To minimize the estimated F1-score across the exon classes, we define:

$$\text{F1-loss} = \sum_{i \text{ is exon class}} (1 - \widehat{\text{F1}}_i),$$

where the sum ranges over three exon classes for the reading frames and eight exon classes for exon border positions.

#### 3.9 HMM

**3.9.1 Distribution**

Let the set of hidden states be $Q = \{1, \ldots, K\}$, let the random variable $Y \in Q^T$ be a state sequence, let $S \in \{\texttt{A}, \texttt{C}, \texttt{G}, \texttt{T}, \texttt{N}\}^T$ be the input DNA sequence, and $X \in [0,1]^{T \times K}$ be the embedding sequence computed from the deep learning layers preceding the HMM. Let $S^{(L)}$ denote the sequence of overlapping, left-adjusted triplets, i.e. $S_j^{(L)} = S[j : j+2]$. Let $S^{(R)}$ denote the overlapping, right adjusted triplets, i.e. $S_j^{(R)} = S[j-2 : j]$. Then the HMM layer defines a joint distribution on the hidden state sequence $Y$ and the layer inputs:

$$P(Y, X, S^{(L)}, S^{(R)}) = \prod_{j=1}^{T} a(Y_{j-1}, Y_j) \cdot b(Y_j, X[j,:]) \cdot c(Y_j, S_j^{(L)}) \cdot d(Y_j, S_j^{(R)}). \quad (1)$$

Here, $a(Y_{j-1}, Y_j)$ is the transition probability, the emission probabilities of the seq2seq embedding are

$$b(Y_j, X[j,:]) = X[j,:]\left(\frac{\alpha}{K-1} + \left(1 - \frac{K-2}{K-1}\alpha\right) e_{Y_j}\right), \tag{2}$$

where $e_k$ is the $k$-th unit column vector ($k \in Q$) and $X[j,:]$ is the $j$-th row of $X$, i.e. the embedding of the deep learning model computed for sequence position $j$. $c(Y_j, S_j^{(L)})$ and $d(Y_j, S_j^{(R)})$ are emission probabilities of triplets starting or ending with $S_j$, respectively. We assume that the emissions of $X_j, S_j^{(L)}, S_j^{(R)}$ are conditionally independent.

###### 3.9.2 Biological Constraints

The HMM layer in Tiberius enforces specific biological patterns at exon border states (see Figure 2). The following constraints are implemented for these states:

- START state: `ATG`
- STOP state: `TAG`, `TAA`, or `TGA`, with emission probabilities 0.34, 0.33, and 0.33, respectively.
- DSS-* states: `GT` or `GC` for start of an intron with emission probabilities 0.99 and 0.01, respectively.
- ASS-* states: `AG` for end of introns.

###### 3.9.3 Parameters

During the fine-tuning, the Tiberius model was fine-tuned end-to-end including the HMM. The HMM itself has only 23 transition parameters, mainly controlling the length distributions of intergenic, intron and exon regions and one parameter for the emission distribution. These parameters were set manually and frozen.

The transition probabilities were chosen based on three expected region lengths $L_{ir} = 10,000$, $L_i = 4,500$ and $L_e = 200$. For $x \in \{ir, i, e\}$, the probability $p_x = \frac{1}{L_x}$ to exit a region was determined as the success probability of a geometric distribution with mean $L_x$. The edges between states of the same region were parameterized with $1 - p_x$. In case of Exon-1 both edges (Exon-1, STOP) and (Exon-1, DSS-2) were parametrerized with $\frac{p_x}{2}$. The emission distribution has only a single parameter $\alpha$. The emission probabilities for all 15 states are set such that state $i$ emits class $i$ with probability $1 - \alpha$ and all other classes with probability $\alpha/14$ (we chose $\alpha = 0.01$). The value $\alpha$ is a smoothing value to account for incorrect inputs. We use a uniform distribution for the initial state $Y_1$.

###### 3.9.4 Parallelization

The forward and backward algorithms are used to compute the posterior state probabilities $\hat{Y}$ and the Viterbi algorithm is used to compute $Y^*$. All three algorithms are parallelized across segments that each sequence of length $T$ is partitioned into defined by breakpoints $1 = j_1 < j_2 < \cdots < j_{k-1} < j_k = T$. Here, the parallel forward algorithm is described. The other two parallel algorithms are analogous.

Parallelization is implemented in two steps. For a sequence $S = s_1, \ldots, s_T$, let $S_{u:v}$ denote the segment $s_u, \ldots, s_v$. First, compute

$$\alpha_{local}(t, q', i, q) := P(Y_{j_t+i} = q, X_{j_t+1:j_t+i} \mid Y_{j_t} = q') \tag{3}$$

in parallel for $t = 1, \ldots, k-1$ and $\forall q, q' \in Q$ and sequentially for $i = 1, \ldots, j_{t+1} - j_t$. Second, from $\alpha_{local}$ compute

$$\alpha_{segment}(t, q) := P(Y_{j_t} = q, X_{1:j_t}) \tag{4}$$

in parallel for $q \in Q$ and sequentially for $t = 1, \ldots, k$.

The recursion formulas are analogous to the sequential forward algorithm:

$$\alpha_{local}(t, q', i, q) = P(X_{j_t+i} | Y_{j_t+i} = q) \sum_{q'' \in Q} P(Y_{j_t+i} = q | Y_{j_t+i-1} = q'') \alpha_{local}(t, q', i-1, q'') \tag{5}$$

for $i > 1$ with the initial condition $\alpha(t, q', 1, q) = P(s_{j_t+1} | Y_{j_t+1} = q) P(Y_{j_t+1} = q | Y_{j_t} = q')$ and

$$\alpha_{segment}(t, q) = \sum_{q' \in Q} \alpha_{local}(t-1, q', j_t - j_{t-1}, q) \alpha_{segment}(t-1, q') \tag{6}$$

for $i > 1$ with the initial condition $\alpha_{segment}(1, q) = P(X_1 | Y_1 = q) P(Y_1 = q)$.

The parallel Viterbi algorithm additionally requires a parallel backtracking operation. To decode the Viterbi sequence, we first compute the optimal states at the breakpoints using $\gamma_{segment}(t, q)$ starting with $t = T$ and thereafter compute all local Viterbi paths using $\gamma_{local}$ in parallel once the right borders of all segments are known. Both passes of this 2-stage backtracking algorithm are similar to the backtracking pass of the sequential Viterbi algorithm.

##### 3.10 Inference of Gene Structures

###### 3.10.1 Merging of Overlapping Tiles

During the inference of gene structures by Tiberius, if the border predictions of adjacent tiles at positions $i$ and $i + 1$ do not match, a reprediction is made on the concatenation of the input sequences of both tiles. Afterwards, the Viterbi parses $h_i$, $h_{i+1}$ of the smaller tiles and the parse $h_{i,i+1}$ of the concatenated prediction have to be merged (see Inference of Gene Structures). This is achieved by taking all complete gene structures that end before the central position of the i-th tile and all that start after this position from the i+1-th tile. For gene structures that overlap the central point, the one with the longest coding sequence is selected. Using the same logic, the gene structures of the first merge and those from $h_{i+1}$ are merged at the central position of $h_{i+1}$.
